# Distinct Working Memory for Near and Far in a T-Maze Delayed Alternation Task

**DOI:** 10.64898/2026.07.30.741937

**Authors:** Masatoshi Takita, Yukio Ichitani

## Abstract

Working memory has been considered a delay-dependent retention system, but variability in delay duration across species and tasks suggests that task-intrinsic factors may also contribute. Here, we investigated the effects of delay duration (75 s vs. 150 s) and task distance (0 m vs. 2 m) on working memory using a T-maze delayed alternation task with a movable home-cage apparatus in rats. At the 150-s delay, accuracy was 9% lower at 0 m than at 2 m (75% vs. 84%), with a similar numerical tendency observed at the 75-s delay (4% lower). A two-way repeated measures ANOVA revealed significant main effects of both task distance and delay duration, with no significant interaction (p = 0.217). Post hoc pairwise comparisons indicated that accuracy in the 0-m condition was significantly lower at 150 s than at 75 s (adjusted p = 0.038). At the 150-s delay, accuracy was also significantly lower in the 0-m condition than in the 2-m condition (adjusted p = 0.019). These results suggest that working memory retention is influenced not only by temporal constraints but also by task-intrinsic factors such as task distance, with the observed effects appearing independent and consistent with a two-factor framework.

## Introduction

Working memory is a cognitive system responsible for the temporary maintenance and manipulation of information over short periods of time (Baddeley & Hitch, 1974). Over the past five decades, working memory has been shown to contribute to a wide range of cognitive functions, including behavioral control and decision-making. Accordingly, various theoretical models have been proposed to explain its mechanisms (e.g., Hitch et al., 2025). Given the limited processing capacity of working memory, investigating the effects of its decline or overload on situation awareness may help clarify the mechanisms underlying human error and contribute to its prevention (Reason, 2000).

Working memory is suggested to share common neurobiological features across primate and rodent species. It is thought to depend on the prefrontal cortex and dopaminergic transmission, and similar neural mechanisms have been proposed in humans (D’Esposito & Postle, 2015). In humans and non-human primates, working memory has typically been studied over delays ranging from seconds to minutes (e.g., Baddeley, 2012). However, in terms of memory precision, studies of human visual working memory have shown that precision declines over delays of several seconds (e.g., Rademaker et al., 2018; Takita et al., 2020). In rodents, working memory has typically been examined using maze tasks, such as the radial maze, with delays ranging from several minutes to several hours (Olton & Samuelson, 1976). At shorter time scales, studies in rats have also examined working memory performance using delayed alternation tasks with second-scale delays in operant conditioning paradigms (Izaki et al., 2008).

Recent reviews suggest that factors beyond the mere passage of time contribute to working memory performance (e.g., Lemaire & Portrat, 2018; Oberauer et al., 2018). Together, these observations raise the possibility that task-intrinsic factors also influence working memory performance.

We focused on one such factor: the spatial extent of the task environment. Whereas operant conditioning tasks are typically performed within restricted spaces, maze tasks require navigation through larger environments. We therefore tested whether this difference affects working memory in rats by manipulating environmental extent in a T-maze delayed alternation task using a movable home cage.

## Results

Figure 1 shows the T-maze with a movable home cage used in the present experiment. Delayed alternation performance was evaluated in six animals according to the pseudo-random schedule shown in Figure 2 under four experimental conditions defined by task distance (0 or 2 m) and delay duration (75 or 150 s) (Fig. 3).

**Figure 1.**
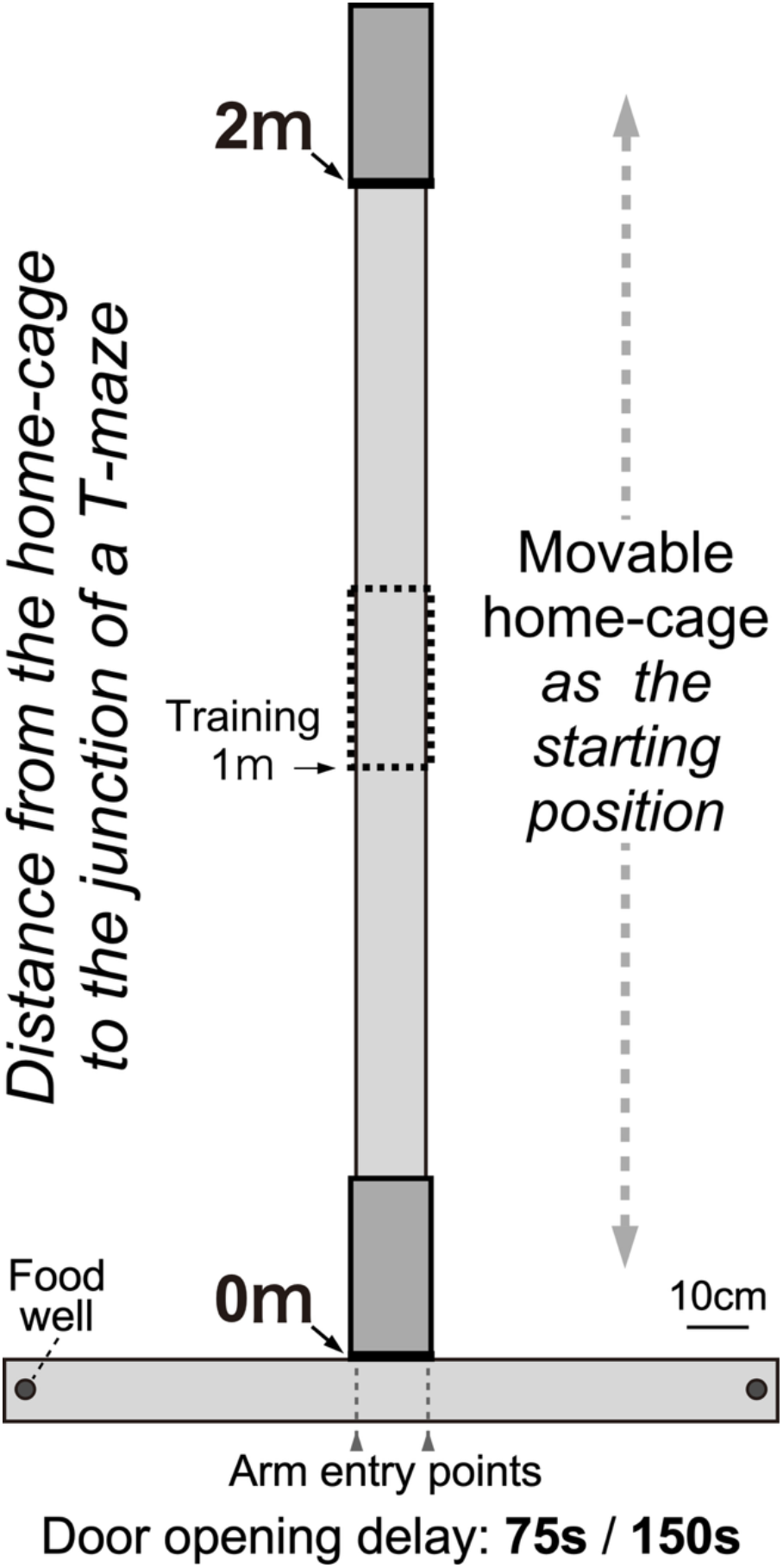
Schematic diagram of a T-maze delayed alternation task for rats. Animals were trained with a movable home cage, with the exit positioned 1 m from the t-junction. Dashed vertical lines indicate the predefined arm entry points. During test trials, the home cage starting position (i.e., maze length) was varied between 0 and 2 m according to a quasi-random schedule (see fig. 2).

**Figure 2.**
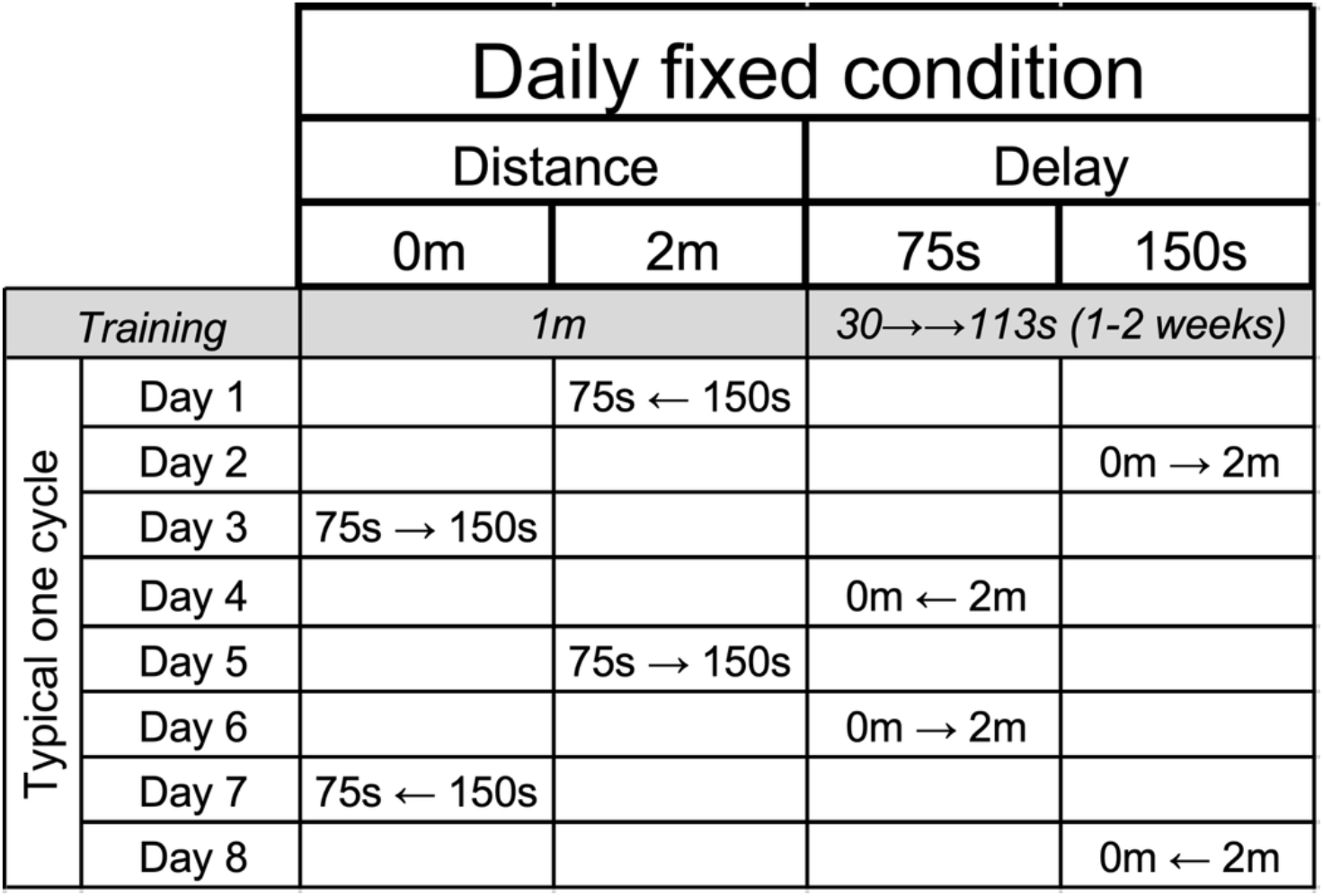
Example of a quasi-random schedule used in the test phase. After training, the four combinations of task distance (0 or 2 m) and delay duration (75 or 150 s) were presented in an 8-day cycle. The conditions were arranged such that consecutive conditions differed in both task distance and delay duration. Each day consisted of two conditions (first and second halves), and arrows (→, ←) indicate transitions between them. Each daily session began with a free-choice trial, in which rewards were placed in both goal wells, followed by delayed alternation trials until rats completed 20 correct trials.

**Figure 3.**
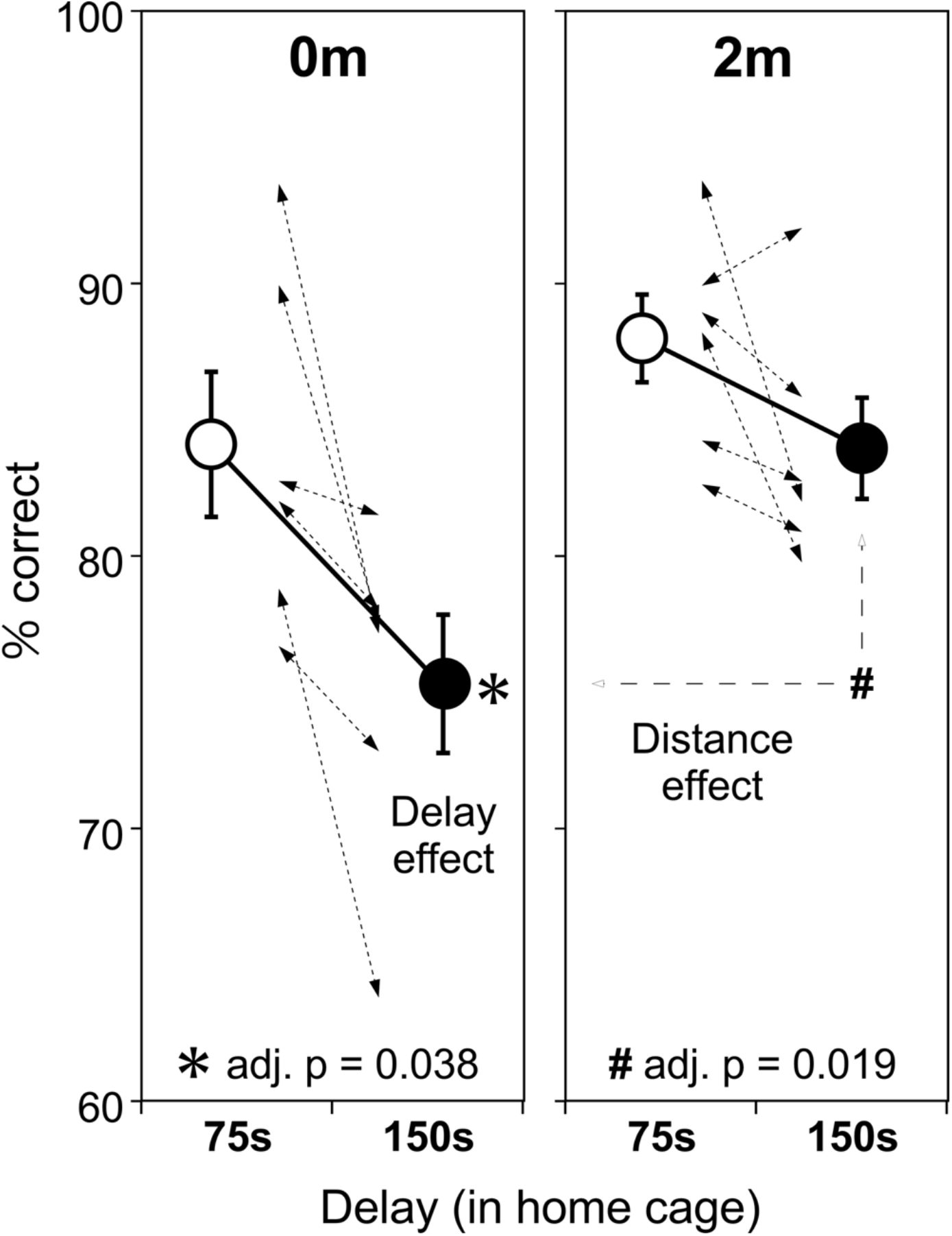
Effects of task distance and delay duration on delayed alternation performance. Delayed alternation accuracy was measured under four conditions defined by task distance (0 m, left panel; 2 m, right panel) and delay duration (75 s, open circles; 150 s, closed circles). Data are presented as mean ± SE (n = 6). Dotted lines illustrate individual changes in performance across delay durations. Two-way repeated-measures ANOVA revealed significant main effects of task distance and delay duration, with no significant interaction. Bonferroni/Dunn-adjusted post hoc pairwise comparisons were conducted to further characterize these effects. These comparisons revealed a significant delay effect in the 0-m condition (adjusted p = 0.038) and a significant distance effect at the 150-s delay (adjusted p = 0.019). * indicates a significant difference between delay durations in the 0-m condition, whereas # indicates a significant difference between task distances at the 150-s delay.

A two-way repeated-measures ANOVA with task distance (0 m vs. 2 m) and delay duration (75 s vs. 150 s) as within-subject factors revealed a significant main effect of task distance (F(1,5) = 17.23, p = 0.0089, η^2^_p_ = 0.71). Accuracy was higher in the 2 m condition than in the 0 m condition. A significant main effect of delay duration was also observed (F(1,5) = 13.95, p = 0.0135, η^2^_p_ = 0.72), with lower accuracy after the 150 s delay than after the 75 s delay. These results indicate that both task distance and delay duration had substantial effects on task performance.

The interaction between task distance and delay duration was not significant (F(1,5) = 2.00, p = 0.2167, η^2^_p_ = 0.38), indicating that the effect of delay duration did not statistically differ between the two task distance conditions. To further characterize the observed main effects, exploratory post hoc comparisons were conducted using the Bonferroni/Dunn correction.

In the 0 m condition, accuracy decreased significantly from 83.8% at 75 s to 75.0% at 150 s (adjusted p = 0.038). In contrast, accuracy in the 2 m condition showed a smaller numerical decrease from 87.9% at 75 s to 83.8% at 150 s, which did not reach statistical significance (adjusted p > 0.1). At the 150 s delay, accuracy was significantly lower in the 0 m condition than in the 2 m condition (adjusted p = 0.019). At the 75 s delay, accuracy was numerically lower in the 0 m condition than in the 2 m condition, although this difference was not statistically significant (adjusted p > 0.2).

Given the small sample size, the estimated effect sizes should be interpreted cautiously because they may be unstable. Nevertheless, the consistent direction of the effects and the large partial eta-squared values suggest that task distance and delay duration may each contribute to behavioral performance.

## Discussion

This study examined the effects of task distance (0 and 2 m) and delay duration (75 and 150 s) on rats’ performance in a T-maze delayed alternation task using a movable home-cage apparatus (Fig. 1) under quasi-randomized conditions (Fig. 2). The present findings indicate that, in addition to the effect of delay duration, task distance influenced performance in the delayed alternation task. Accuracy was higher in the 2 m condition than in the 0 m condition. This finding was unexpected because the present study was initially motivated by attempts to shorten the training period in the movable home-cage paradigm, based on the assumption that reducing the distance between the home cage and the choice point might facilitate performance. Instead, the present findings suggest that task distance may influence how animals engage with the task and perform the required choice behavior. This distance effect may reflect differences in behavioral strategies or response selection processes associated with different maze distances (Fig. 3). The delay durations used in this study (75 s and 150 s) were selected with reference to the delay durations presented in Fig. 4 of Dudchenko (2001). Previous work has shown that working memory retention in a T-maze can be maintained even with delays exceeding one hour (Lett, 1975).

To our knowledge, few studies have directly examined the effect of task distance on working memory using T-maze paradigms with controlled distance manipulations. In an auditory discrimination task, no performance differences were observed across T-maze lengths ranging from 20 to 80 cm (Holleman et al., 2019). Studies using radial-arm maze tasks reported distance-related effects, with reduced performance in a short-arm condition (37 cm) and improved performance in a long-arm condition (80 cm), which were interpreted as reflecting changes in choice criteria rather than alterations in working memory capacity (Brown, 1990; Brown & Huggins, 1993). Together, these findings suggest that task distance may influence behavioral strategies or response selection processes underlying working memory performance rather than memory capacity itself.

Considering parameters reported in previous studies (120 s for maze-based tasks [Suenaga et al., 2008] and 16 s for operant tasks [Izaki et al., 2008]) some commonly used human and non-human primate working memory paradigms may share the characteristic of limited movement requirements with the 0 m condition because they rely primarily on stationary responding rather than movement through an extended environment. Therefore, differences in task structure and response requirements should be considered when comparing working memory performance across experimental paradigms.

One possible interpretation is that differences in task distance alter the mode of working memory operation rather than simply increasing or decreasing mnemonic load. Under this view, task distance may influence how memory maintenance and action selection are coordinated during task performance, leading to different behavioral outcomes despite similar nominal memory demands. Thus, task distance may represent one type of task-intrinsic factor that shapes working memory processing, raising the possibility that different task structures engage different modes of working memory operation. The neural mechanisms underlying these task-dependent effects remain unclear and require further investigation using electrophysiological and other approaches.

Overall, the present study demonstrates that task distance influences performance in a T-maze delayed alternation task beyond the effects of delay duration. These findings suggest that working memory performance is shaped not only by memory demands but also by task-intrinsic factors that influence how animals engage with and execute the task. The present results provide a behavioral framework for future investigation of the neural basis of task-dependent modes of working memory operation.

## Materials and methods

### Animals

All animal experimental procedures were approved by the Institutional Animal Care and Use Committee of National Institute of Advanced Industrial Science and Technology (approval numbers: LS-00000799, LS-00000968) and were conducted in accordance with the guidelines of the National Institutes of Health (NIH, 1996) and the Science Council of Japan (2006). The data obtained in this study were analyzed at the University of Electro-Communications.

Seven 6-week-old male Sprague–Dawley rats (CLEA, Tokyo, Japan) were purchased and housed individually. One rat was used in a preliminary study to establish the behavioral training protocol and was not included in the final analysis. The remaining six rats were used in the main experiment. After an acclimation period of 2–3 weeks, their body weights ranged from 270 to 317 g at the start of behavioral training. Body weight was maintained at ≥85% of the free-feeding weight (according to CLEA standards). Rats were fed standard laboratory chow (CE-7, CLEA, Tokyo, Japan) with ad libitum access to water. The animal room was maintained at 24°C under a 12 h light/dark cycle.

### Devices and settings

The T-maze was designed based on the apparatus described by Suenaga et al. (2008). Briefly, a T-shaped maze constructed from matte black acrylic (corridor width, wall thickness, and wall height: 12, 1, and 4 cm) was elevated 50 cm above the floor. The maze consisted of a 134-cm crossbar and a 230-cm stem extending from the center of the crossbar. The movable home cage (external dimensions [D × W × H]: 30 × 14 × 30 cm) was equipped with a manually operated cage door (clear opening, 14 × 30 cm), allowing the task distance to be adjusted from 0 m to 2 m (Fig. 1). Except for the central 12 × 12 cm region of the crossbar adjacent to the stem, each left and right arm measured 60 cm in length. Food wells (3 cm in diameter and 1 cm deep) were placed at both ends of the crossbar, with a center-to-center distance of 125 cm.

During the experiment, broadband noise (60–65 dB SPL at the center of the T-maze) was continuously delivered through two speakers mounted near the ceiling. Illumination was provided indirectly by four ceiling-mounted LED lights, maintaining a light intensity of 9–10 lx within the maze corridors.

Each session began when the cage door was opened. Simultaneously, a 12 V DC LED light positioned above the rear of the home cage was turned on via a foot switch, illuminating the interior of the cage (approximately 140 lx). When the rat reached a food well, the light was turned off, and the entrance to the opposite arm was blocked using a gate panel identical in size to the door. After the animal voluntarily returned to the cage, the cage door was closed. Animal behavior was recorded at a standard video frame rate using a pinhole-type CCD camera (sensitivity: 0.5 lx) mounted near the ceiling above the T-junction.

### Training and Testing

Training began with habituation, with the distance between the movable home cage exit and the T-junction set at 1 m. As a reward, 45-mg dustless precision pellets (Bio-Serv, Flemington, NJ, USA; F0021-J) were used. On the first day of training, pellets were placed at approximately equal intervals along the maze and in the food wells (7 pellets initially, then 5). Three pellets were placed at the crossbar section, followed by repeated sessions in which rats freely chose between the left and right wells, after which only one pellet was available in an alternating manner. The total number of pellets provided on the first training day was 21. From the following day, the delay period was gradually increased from 30 s based on each animal’s performance, and training continued until the rats achieved a success rate of 80 % (number of correct trials / total trials) at a delay of 113 s (approximately 1–2 weeks).

In the main experiment, each session was conducted under one of four combinations of task distance (0 m or 2 m) and delay length (75 s or 150 s), presented according to a pseudorandom schedule in which identical or similar conditions did not occur consecutively (Fig. 2). One cycle consisted of one session per day for eight consecutive days, and the data for each animal were averaged across 3–5 cycles.

Each session began with a free-choice trial, in which rewards were placed in both goal wells, followed by 20 delayed alternation trials. After the rat achieved 10 correct responses in the first half condition, the session proceeded to the second half condition and ended after 10 correct responses were achieved in that condition.

When switching between distances (0 m and 2 m), the arm exit was temporarily closed with a gate panel at the moment the animal obtained the reward, and the position of the movable home cage was changed during this interval. Immediately after training or each session, the rats were given one pellet of standard chow (approximately 3 g) and then returned to their home cages.

### Behavioral events

Arm entry was operationally defined as crossing the predefined arm entry point in the arm where the rat ultimately reached the food well (Fig. 1). This criterion was applied uniformly to all trials, including rare instances of vicarious trial-and-error behavior, in which rats briefly entered one arm but reversed before reaching the food well in that arm and subsequently reached the food well in the opposite arm.

## Data analysis

Working memory performance was evaluated based on the percentage of correct responses for each condition, separately for the first and second halves of each daily session (see Fig. 2). Statistical analyses were performed using repeated-measures analysis of variance (ANOVA), followed by post-hoc pairwise comparisons using the Bonferroni/Dunn method (Dunn’s test with Bonferroni adjustment), as implemented in commercial software (StatView J-5; Hulinks Inc., Tokyo, Japan). In all cases, P < 0.05 was considered to indicate statistical significance. All data are expressed as mean ± SE.

## Acknowledges

We express our deepest gratitude to the late Dr. Yoshinori Izaki (St. Marianna University School of Medicine, who passed away on June 20, 2011) engaged in valuable discussions with M.T. on working memory tasks in operant and maze paradigms. We thank Drs. Hiroshi Yokoi and Yoshiko Yabuki (The University of Electro-Communications) for their valuable discussions, and Ms. Atsuko Yamashita (National Institute of Advanced Industrial Science and Technology [AIST]) for her technical assistance. This work was supported by JSPS KAKENHI (15K04202 and 18K03196 to M.T.), and in part by an AIST grant for neurorehabilitation research (M.T.). Part of this work was conducted while M.T. was affiliated with AIST.

